# Feather corticosterone is not associated with feather growth rate or quality across tropical and temperate passerines, but possibly linked to elevation

**DOI:** 10.64898/2026.01.09.698619

**Authors:** Ondřej Kauzál, Oldřich Tomášek, Kryštof Horák, Marie Kotasová-Adámková, Tereza Kauzálová, Jacques E. Chi, Francis T. Mani, Zuzana Hochmanová, Zdeněk Šimek, Tomáš Albrecht

## Abstract

Moult is a challenging stage in the life of birds. Small passerines replace plumage, substantial part of their body mass, annually to preserve its vital functions. Surprisingly, most species tend to downregulate corticosterone, major mediator of energy metabolism and stress response, during this energetically demanding period. Experimental studies suggest corticosterone has detrimental effect on feather quality and, within species, slows feather growth rate (FGR), prolonging the period when plumage functions are compromised. However, how exactly corticosterone affects moult in natural populations or on macroecological scale is not yet fully understood. In this study, we analysed corticosterone deposited in tail feathers (fCORT), a corticosterone cumulative measure across multiple days relevant to the moult, on an unprecedented sample of 87 passerine species from Europe and Afrotropics. Our data showed moderate between and high within-species repeatability, strengthening the usefulness of fCORT as a tress record across multiple species. We did not find any association of fCORT with either FGR or fault bar occurrence. Unlike circulating corticosterone, fCORT did not differ between latitudes. However, our data suggested higher fCORT in high elevation tropical species. More research is needed to understand how birds regulate corticosterone during moult and how it affects moult in natural populations.

## Introduction

In vertebrates, growth, physiological and behavioural processes are regulated by hormones. They mediate stress response, trigger major life history events like breeding, migration or, in case of birds, the annual moult or feather replacement (Sapolsky et al. 2000, Dawson 2008). Among the most studied hormones is corticosterone (CORT), the primary avian glucocorticoid regulating energy balance and acute stress response (Wingfield et al. 1998, Sapolsky et al. 2000). Circulating levels of CORT vary wildly among bird species but they seem to be closely associated with the pace of life – tropical species, which typically show lower basal metabolic rates, reduced annual mortality and slow pace of reproduction, generally exhibit lower baseline CORT than their fast-living temperate counterparts (Hau et al. 2010, Kauzal et al. 2025). Within species, circulating CORT also changes dramatically across distinct life-history stages, including breeding, migration or moult (Romero 2002, Bauer & Watts 2021).

Moult is a key life-history stage during which passerines replace all their feathers at least once a year to counteract their natural wear (Jenni & Winkler 2020a). Moulting represents a major energetic challenge, because plumage, predominantly made of keratin (Murphy 1996), can make up to 40 % of the bird’s dry mass (Ginn & Melville 1983). Moreover, passerines typically complete moult within a relatively short period (40-70 days in most temperate Passerines; Ginn & Melville 1983), which necessitates a substantial increase in metabolic rate (Lindström et al. 1993). Although elevated CORT can facilitate energy acquisition (Piersma et al. 2000, Eikenaar et al. 2014), circulating CORT is often downregulated, rather than upregulated, during moult (Romero 2002), presumably because elevated CORT can directly compromise feather growth and quality by redirecting energy towards mechanisms related to self-maintenance (Sapolsky et al. 2000). Indeed, experimentally elevated CORT during moult has detrimental effect on growing feathers, causing structural weakening and slowing feather growth (Romero et al. 2005, Strochlic & Romero 2008, DesRochers et al. 2009, Lattin et al. 2011, Jenni-Eiermann et al. 2015). This is important, because reduced feather quality can impair flight performance, thereby reducing predator evasion and foraging success and ultimately individual fitness (Echeverry-Galvis & Hau 2013, Hedenström 2023). Slower feather growth could also directly prolong the moulting period. That would increasing time when individuals experience wing gaps, and therefore reduced flight capability, while also potentially reducing the plumage thermal insulation and elevating energetic demands even further (Jenni & Winkler 2020a, Hedenström 2023).

Measuring CORT in feathers (fCORT) is an elegant and recently very popular method to measure the level of circulating CORT during moult, without the need of sampling individuals during time when most birds become very secretive (Jenni & Winkler 2020a). CORT is deposited in feathers during their growth (when they are vascularized; Jenni-Eiermann et al. 2015) and once deposited, fCORT is stable and can be measured years after the feather had finished growing (Bortolotti et al. 2009, Beattie & Romero 2023). This offers a unique opportunity to sample CORT directly relevant to the moulting period (Harris et al. 2016, Romero & Fairhurst 2016). fCORT is useful as an integrated history of circulating CORT over an extended period of time (up to several weeks in case of flight feathers) and likely reflects both baseline levels and stress-induced peaks (Romero & Fairhurst 2016).

This “stress record” indeed reflects experimentally elevated circulating levels of CORT (Romero et al. 2005, Strochlic & Romero 2008, DesRochers et al. 2009, Lattin et al. 2011, Jenni-Eiermann et al. 2015) and correlates with developmental stress (Will et al. 2014) or environmental conditions (Legagneux et al. 2013, Treen et al. 2015) in natural populations. Negative association of fCORT with FGR has been observed both in experimental studies (Romero et al. 2005, Jenni-Eiermann et al. 2015) as well as in some natural populations (Adámková et al. 2019, Sonnenberg et al. 2024, Nikolaou et al. 2025) and variation in fCORT has been also linked with traits not directly associated with moult, including survival (Koren et al. 2012, Lind et al. 2020) highlighting its broader utility as an integrative biomarker. However, to our knowledge, no study has yet tested whether fCORT shows macroecological variation (e.g., tropical vs. temperate zone taxa, high vs. low elevations) analogous to that reported for circulating CORT (Hau et al. 2010, Kauzal et al. 2025) or other POLS (such as basal metabolic rate or blood glucose (Wiersma et al. 2007, Tomášek et al. 2022).

Here, we present, to the best of our knowledge, the first comparative analysis of fCORT across multiple passerine species sampled across various habitats in the Palearctic and Afrotropical realms. We collected a unique dataset of fCORT concentrations measured with a high-sensitivity method in tail feathers from 352 individuals of 87 species spanning 33 passerine families, complemented by already published data of feather growth rates (FGR) and the occurrence of fault bars (FBO) from the same feathers (Horák et al. 2022). Using this dataset we tested following predictions: (1) within species, fCORT concentrations will be negatively associated with FGR (Jenni-Eiermann et al. 2015, Adámková et al. 2019). Across species, and controlled for potential effect of latitude (Horák et al. 2022), we expect either a similar pattern – fCORT will be negatively associated with FGR, or there will be no consistent association across species as circulating CORT is generally downregulated during moult (Romero 2002) and species could differ in their sensitivity to circulating CORT to avoid its detrimental effects on feather growth (Lattin et al. 2011, Jenni-Eiermann et al. 2015); (2) Because of detrimental effect of CORT on feather quality, we expect FBO to be positively associated with fCORT when controlled for potential effect of latitude (Lattin et al. 2011, Jenni-Eiermann et al. 2015, Horák et al. 2022). We further evaluated whether fCORT covaries with geographic and life history traits that are known to predict circulating plasma CORT. Specifically, we predicted that (3) fCORT will be associated positively with feather mass and body mass (Romero & Fairhurst 2016) and will be either lower in tropical species parallelling patterns in circulating plasma CORT (Hau et al. 2010, Kauzal et al. 2025) or higher in tropical species because of their higher FBO and lower FGR (Horák et al. 2022). (4) Temperate migratory species will show either lower fCORT to facilitate rapid growth of high-quality feathers (Romero et al. 2005, De La Hera et al. 2012) or higher fCORT due to elevated energetic demands during a comparatively shorter moulting season (De La Hera et al. 2009, Kiat et al. 2019) than sedentary species. Finally, we expect (5) fCORT will be higher in tropical high-elevation species, consistent with harsher environment possibly shortening moult period or putting higher energetic demands (Hernández=Téllez et al. 2024, Sonnenberg et al. 2024).

## Material and Methods

### Study species and field sites, feather sample collection

We trapped small passerines using mist nets during regular fieldworks in Czechia (temperate zone site, 48.7-50.1 °N, 13.9-17.2 °E, 170-730 m a.s.l.) and in Cameroon (3.9-6.1 °N, 9.0-10.3 °E, 10-2280 m a.s.l.; Fig S1) in 2013-2018. To cover as ecologically and taxonomically diverse spectrum of passerine species as possible, mist netting was conducted in various habitats – from streams, reedbeds, open shrubland through secondary and primary forest to urban areas. We then selected 352 adult individuals of 87 passerine species (40 temperate and 47 tropical) averaging 4.0 samples/species (*s.d.* = 1.39) for further analysis.

### Field data & molecular sexing

After capture, a feather sample was collected consisting of both second outermost rectrices (i.e. tail feathers “R5”; Fig. S2). Birds were then ringed with a unique metal ring (issued either by the Czech Ringing Centre or SAFRING), weighed and a standard set of morphological measurements were taken. Also, a small blood sample was taken for each bird and stored in 96% ethanol. Whenever possible, birds were aged using plumage criteria (Svensson 1992, Demongin et al. 2016) and sexed using either plumage criteria or by inspecting the development of brood patch or cloacal protuberance. Because brood patch and cloacal protuberance development can be unreliable for sexing in poorly known monomorphic tropical species, all individuals of tropical species were molecularly sexed (for details see Horák et al. 2022).

### Feather growth rate, fault bar occurrence and other life history traits

We compiled life history data for each species from literature and other available sources. For each analysed feather, individual and average species specific FGR, FBO, feather growth time and feather (rhachis) length data were taken from our previous work on an overlapping dataset (Horák et al. 2022), as well as the information about species-specific elevational midpoint (i.e. at what elevation species lives at; Tomášek et al. 2022). The information about the moulting latitude (latitude at which species moults its rectrices) was compiled from Jenni & Winkler (2020b). We then classified birds into three categories based on their breeding and moulting latitude (further referred to as breeding-moulting category, *BMC*): i) temperate zone breeding species moulting in temperate zone, ii) temperate zone breeding species moulting in tropics, iii) species breeding and moulting in tropics (Horák et al. 2022). Migration distance was calculated as latitudinal arch difference (in 1000km) between the centroid of breeding and wintering range (BirdLife International and Handbook of the Birds of the World 2018; for details see Supplementary materials and methods). We did not have body mass for 32 out of 352 individuals, so we calculated average species body mass from our data instead (*n* = 7290). The correlation between individual and average body mass in the final dataset was very high (Pearson’s *r*≈ 0.99, *n* = 320, *p* < 0.001). To test for possible confounding effect of common ancestry, we used a phylogenetic tree which we previously constructed using publicly available DNA sequences (for details see Horák et al. 2022).

### LC-MS/MS analysis

For the analyses of fCORT, one of the sampled feathers (mostly the left one) with its calamus (part of the feather without the vane) clipped off, further on referred to simply as *feather*, was weighted on analytical scales (precision 0.00001 g). Individual feathers were then clipped into smaller pieces and pulverized using a ball mill (Mixer Mill MM 200; Retsch) at 30 Hz for 120 min using 3 mm stainless steel grinding balls. After pulverization, samples were randomized prior analysing in an analytical chemistry laboratory in RECETOX (Brno, Czechia) using liquid chromatography–tandem mass spectrometry (LC-MS/MS) and a newly developed method utilizing hormone derivatization to increase sensitivity and detection selectivity (for details see Bílková et al. 2019).

Out of the 352 samples of fCORT analysed, 38 were under quantification limit (LOQ = 0.83 pg in the final 60 μl extract; Adámková et al. 2019; Table S1). Their distribution in the dataset was random with respect to sex (females: 9.8%; males: 11.6%; χ² ≈ 0.12, *p* ≈ 0.72) but not with respect to BMC (temperate–temperate: 12.6%; temperate–tropical: 23.9%; tropical–tropical: 6.1%; χ² ≈ 12.7, *p* ≈ 0.002).

To obtain within-individual repeatability of fCORT we also analysed the second tail feather for 46 individuals of 25 species from the original dataset. These data were used only in the repeatability estimation analysis.

### Statistical analysis

We expressed final values of fCORT as ln-transformed absolute hormone amounts (fCORT_abs_ [pg]), rather than concentrations. We further include ln-transformed average body mass, feather length and growth time in all models as covariates. This approach allowed us to estimate how fCORT scales with body and feather size and growth time, rather than constraining their effects a priori (i.e. forcing the slope to 1).

For each individual, we calculated the standardized difference of individual FGR from species-specific average (z-score) from the ln-transformed values (further referred to as individual feather growth deviation, *IGRD*). For elevation models, we calculated the difference of the elevation of capture from the species-specific elevational midpoint. This roughly represents how far from the optimal species elevation was the individual bird caught. Following variables were coded as dummy variables: sex (0: male, 1: female) and breeding and moulting latitude (0: temperate, 1: tropical).

Data were analysed using the Bayesian phylogenetic mixed models based on the Hamiltonian Monte Carlo algorithm using the brms package v. 2.22.0 (Bürkner 2017) in R v. 4.4.2 (R Core Team 2023). Default priors defined in the brms package were used and the models were run in 8 chains, each with 8,000 iterations, warm-up of 4,000 and thinning of 1. For detailed information about brms models, their validation and comparison see (Kauzal et al. 2025). All results are presented as posterior means (*b*) with quantile-based 95% credible intervals (CrI95). The support for an effect is considered substantial when CrI95 does not contain zero and weak when the probability of direction (*pd*) of the parameter being different than zero is larger than 95% (i.e., >95% of the posterior mass was on one side of zero), which corresponds approximately to a two-sided frequentist *p-value* < 0.10 (Makowski et al., 2019).

For models where fCORT was the response variable, we treated the values for fCORT below the quantification limit (“left-censored data”) in four different ways: i) using the native *cens* function of the brms package for treating left-censored data, ii) substituting LOQ with a fixed value of LOQ/2 (0.415 pg), iii) using a random value below LOQ from an estimated distribution and iv) excluding the values below LOQ from the models (for details see Supplementary Materials and methods). All methods of imputation performed similarly and produced very similar results (Table S2). In the main text, we present the models with values below LOQ imputed using the *cens* function over other methods of imputing data. This method is advantageous as it accounts for uncertainty of the imputed values (Bürkner 2017). For the FBO and FGR models where fCORT was a predictor, we were unable to use the *cens* function to impute the data. Instead, we substituted the values below LOQ with random values from estimated distribution, or as LOQ/2, or removed them completely and compared the models. All models performed similarly (Table S2) and we report only models using random values for LOQ in the main text.

First, we run an exploratory set of simple phylogenetic regression models to assess how fCORT scales with body and feather mass, feather length and growth time and how body mass scales with feather mass and length (Table S3). All further analyses were then controlled for the species ln-transformed average body mass, feather length and growth time, except for FGR models which were controlled only for species average body mass, because FGR is calculated as feather length/growth time. Before running the main models, all predictors were also tested individually in models hereafter referred to as “adjusted single effect” (ASE) models to test their overall effects. All the ASE models were controlled for the species average body mass, feather length and growth time (Table S4). In all models, we have included LC-MS/MS run ID, species ID and phylogeny as random effects in order to control for potential variation among runs, non-independence of multiple observations per species, and shared ancestry, respectively.

### Ethical statement

The study was conducted in accordance with the Guidelines for Animal Care and Treatment of the European Union and was approved by the Animal Welfare Committee of the Czech Academy of Sciences under protocol numbers 041/2011 and 09/2015 and the Cameroon Ministry of Research and Innovation under the research permit numbers 0021/MINRESI/B00/C00/C10/C14 and 0412/PRS/MINFOF/SG/DFAP/SDVEF/SC.

## Results

We analysed 352 samples of 87 passerine species (40 temperate and 47 tropical). We were able to obtain fCORT for 314 out of 352 analysed samples (Table S1). Repeatability of fCORT_abs_ within species was moderate (≈0.30; CrI95: 0.19, 0.43) but high within individuals (≈0.69; CrI95: 0.44, 0.85; *n* = 46 individuals, 25 species). Across species, on log–log scale, fCORT_abs_ scaled with species body mass with a slope of 0.78, with feather mass with a slope of 0.58, and with feather length with a slope of 0.94. Feather mass scaled almost linearly on log–log scale with body mass (slope ≈ 0.99), whereas feather length did not (slope ≈ 0.29; for CrI and marginal R^2^ of the slopes see Table S3). Results of ASE models are given in Table S4.

First, we tested whether FGR or FBO was associated with fCORT. FGR was positively associated with average species-specific body mass and was lower in tropical species. There was no association with species specific fCORT_abs_ controlled for average body mass but the model suggested a very slight negative association with the individual deviation from the average species-specific. However, the probability of direction was below 90%, therefore we rejected it as being unsupported by our data (Table 1). The model with FBO as response variable showed no association with fCORT_abs_ controlled for average body mass and feather length or sex, but as expected showed higher FBO in tropical breeders (Table 2).

**Table 1:** Summary of the main model testing the effect of breeding and moulting latitude (BMC), sex and feather corticosterone (fCORT) on feather growth rate. Apart from phylogeny and run ID, the models were controlled for species specific average body mass, i.e. representing the differences in fCORT for same-sized birds. Values with CrI95 not containing zero are highlighted in **bold** and regarded as significant support for an effect. Weakly supported trends in models (*pd* between 95% and 97.5%) are highlighted in *italic*.

| predictor | *b* | LCrI | UCrI |
| --- | --- | --- | --- |
| BMC: temperate–tropical | 0.14 | −0.10 | 0.38 |
| **BMC: tropical–tropical** | **−0.35** | **−0.51** | **−0.19** |
| sex (female) | 0.00 | −0.06 | 0.06 |
| ln species specific fCORT<sub>abs</sub> | 0.01 | −0.08 | 0.11 |
| fCORT<sub>abs</sub> deviation from average | −0.02 | −0.05 | 0.01 |
| **ln species specific body mass** | **0.49** | **0.32** | **0.66** |
| effect of phylogeny | 0.76 | 0.61 | 0.84 |
| Bayesian conditional R<sup>2</sup> | 0.81 | 0.78 | 0.83 |
| Bayesian marginal R<sup>2</sup> | 0.39 | 0.27 | 0.51 |

**Table 2:** Model summary of the model testing the association between fault bar occurrence and breeding and moulting latitude (BMC), sex and feather corticosterone (fCORT) using values imputed from the Gaussian distribution. Apart of phylogeny and run id, model was controlled for species specific body mass, feather length and growth time. Values with CrI95 not containing zero are highlighted in **bold** and regarded as significant support for an effect.

| **predictor** | ***b*** | **LCrI** | **UCrI** |
| --- | --- | --- | --- |
| BMC: temperate–tropical | 0.44 | −1.85 | 2.52 |
| **BMC: tropical–tropical** | **2.61** | **1.38** | **4.11** |
| sex (female) | 0.11 | −0.58 | 0.81 |
| ln species specific fCORT<sub>abs</sub> | −0.08 | −0.71 | 0.57 |
| fCORT<sub>abs</sub> deviation from average | −0.16 | −0.56 | 0.24 |
| ln species specific body mass | 0.68 | −0.46 | 1.86 |
| ln feather length | −1.01 | −3.35 | 1.14 |
| ln growth time | 2.18 | −0.63 | 5.20 |
| effect of phylogeny | 0.12 | 0.00 | 0.46 |
| Bayesian conditional R<sup>2</sup> | 0.21 | 0.12 | 0.31 |
| Bayesian marginal R<sup>2</sup> | 0.13 | 0.04 | 0.23 |

Then, we tested whether fCORT_abs_ changes with breeding and moulting latitude (BMC) and if this variation is sex-dependent using a model with an interaction between BMC and sex and a model without this interaction (Table S5). In both models fCORT_abs_ scaled positively with average species-specific body mass (and therefore also with the feather mass), but after accounting for this body mass effect, both feather length or growth time showed no clear association with fCORT_abs_. The model with interaction indicated potentially higher fCORT_abs_ for females in temperate breeding and moulting species compared to species moulting in tropics (*pd* = 96%). The model without interaction suggested a trend towards lower fCORT_abs_ in temperate breeding birds moulting in tropics compared to both temperate breeding and moulting birds and tropical birds, but the probability of this effect not being zero was only 90% (Table S5). Model comparison did not clearly favour either of the models, nor was there substantial evidence that either model differed from models without BMC or sex (Table S6). Therefore, we treat BMC and sex as not being considerable predictors of fCORT.

To test the effect of migration distance we then constructed a similar model with migration distance as a predictor variable on a subset of temperate species (*n* = 171 ind. of 40 species). This model suggested a trend for fCORT_abs_ to slightly decrease with migration distance, but since the probability of direction was only 94%, we rejected it as not supported (Table S7).

Finally, for a subset of tropical species (*n* = 172 ind. of 46 species) we also tested whether fCORT_abs_ is associated with the species-specific elevation midpoint and the individual difference from this midpoint. There was some evidence that tropical species living at higher elevations had higher fCORT (CrI95 slightly overlapping zero, *pd* = 97%, Fig. 1), while the individual deviance from the elevational midpoint showed no clear association with fCORT (Table 3).

**Fig. 1:**
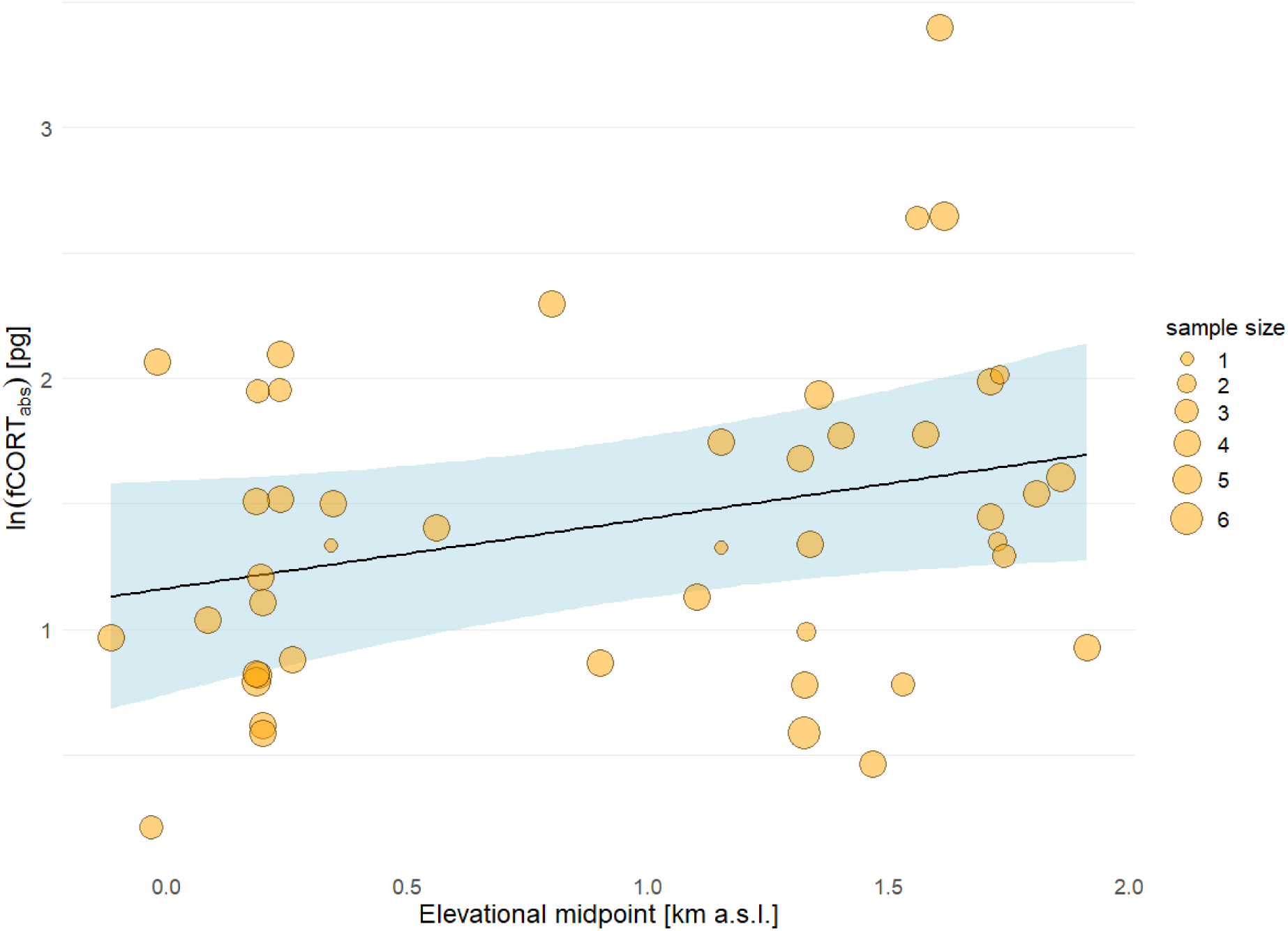
Relationship between species specific elevational midpoint and feather corticosterone (fCORT) for tropical species (*n* = 172 ind. of 46 species; for details of the model see Table 3). Circles represent the average values of fCORT_abs_ per species and their size correspond to their sample size.

**Table 3:** Summary of a model testing the effect of elevation (species specific and individual difference) on feather corticosterone (fCORT) in a subset of tropical species (*n* = 172 ind. of 46 species). Apart from phylogeny and run ID, the model was controlled for species specific average body mass, feather length and growth time, i.e. representing the differences in fCORT for same sized bird with same sized feather per one day. Values with CrI95 not containing zero are highlighted in **bold** and regarded as significant support for an effect. Weakly supported trends in models (*pd* between 95% and 97.5%) are highlighted in *italic*.

| **predictor** | ***b*** | **LCrI** | **UCrI** |
| --- | --- | --- | --- |
| sex (female) | 0.04 | -0.22 | 0.31 |
| *species elevation midpoint* | *0.28* | *-0.01* | *0.58* |
| diff. from species elevation midpoint | -0.15 | -0.49 | 0.20 |
| **ln species specific body mass** | **0.94** | **0.41** | **1.50** |
| ln feather length | -0.35 | -1.43 | 0.71 |
| ln growth time | 0.87 | -0.38 | 2.12 |
| effect of phylogeny | 0.22 | 0.00 | 0.55 |
| Bayesian conditional $R^2$ | 0.47 | 0.37 | 0.57 |
| Bayesian marginal $R^2$ | 0.30 | 0.16 | 0.42 |

## Discussion

In this study, we present the results of a comparative study of corticosterone deposited in feathers during their growth (fCORT) across 87 passerine species breeding in temperate and tropical zones while controlling for key covariates. Using Bayesian modelling we quantified variation both within as well as across species in our dataset. To the best of our knowledge, this is the first study using data collected with a standardised protocol and measured in a single laboratory, examining both interspecific and intraspecific variation in fCORT. We analysed the samples with recently developed method utilizing derivatization of hormones for increased sensitivity in complicated matrices such as feathers (Bílková et al. 2019) and found moderate within-species repeatability (≈0.30), comparable to estimates to other physiological traits like circulating CORT (Ouyang et al. 2011, Kauzal et al. 2025) or blood glucose (Tomášek et al. 2022). We have also estimated the within-individual repeatability by analysing two outermost tail feathers (left and right), which normally grow at the same time and should therefore reflect the same levels of circulating CORT (Jenni & Winkler 2020a). The within individual repeatability was high (≈0.69) and comparable to studies using same feather tract for repeatability estimation (e.g. Adámková et al. 2019), but much higher than in studies using different feather tracts (e.g. Harris et al. 2016). If anything, the inherent and unavoidable laboratory error should lower the repeatability estimate. Therefore, our high individual repeatability further supports that fCORT measurements reliably capture an individual “stress-record” over a certain period of time.

Unlike many studies that report fCORT as concentration (either per feather mass or length – to standardize for the size of the sample; e.g. Bortolotti et al. 2008, Lattin et al. 2011, Koren et al. 2012, Treen et al. 2015), we report the measured fCORT as absolute values and we control in the models for average body mass (and therefore for tightly correlated feather mass), feather length and feather growth time (Bortolotti et al. 2008, Jenni-Eiermann et al. 2015). Expressing fCORT as concentrations is equivalent to setting the slope between measured CORT and feather length (or mass) to 1 and can have some statistical drawbacks including “spuriously significant results and/or biased effect size estimates” (Nakagawa et al., 2017). While such approach might be appropriate in single species studies where the variation in feather length/mass between individuals is small, variation in feather mass in our dataset was considerable (0.9–43.4 mg) and, on the original scale, fCORT did not scale linearly with neither feather mass nor body mass (i.e. slope ≠ 1; Table S3, Fig. S3), similarly to other studies (Lattin et al. 2011, Romero & Fairhurst 2016, Freeman & Newman 2018). This nonlinear allometry might be both natural due to the mass-dilution of CORT and structural differences in keratin density (Bortolotti et al. 2008, Romero & Fairhurst 2016), or due to the technical limitations of measuring CORT in small feather samples (Lattin et al. 2011, Freeman & Newman 2018). Expressing fCORT per feather length over feather mass might be a more suitable option (as suggested by e.g. Bortolotti et al. 2008, Jenni-Eiermann et al. 2015) and our data are broadly consistent with this view: fCORT scaled with feather length with a slope close to 1. However, we used both variables in main models mainly to control for the actual size of the feather sample rather than to interpret their slopes biologically.

Neither feather growth rate (FGR; Table 1), nor the fault bar occurrence (FBO; Table 2) were clearly associated with fCORT, either within or across species. This is rather surprising considering that on an individual scale, the detrimental effect of artificially induced CORT on FGR reported by experimental studies using implants is considerable (e.g. Romero et al. 2005, Jenni-Eiermann et al. 2015). However, in a study using more natural stressors during the period of moult, the observed increase in CORT due to stress was much smaller, suggesting that in nature, birds tend to avoid high CORT during moult even when stressed and CORT implants may not represent well the natural circulating stress levels (Strochlic & Romero 2008). That could explain why we observed only a very weak trend in our dataset derived from natural populations (but see e.g. Adámková et al. 2019). It is possible that other factors such as nutrition and condition or heritability might have bigger impact on FGR variation in the natural populations (Strochlic & Romero 2008, De La Hera et al. 2022). Also, on a macroecological scale, species might have simply evolved different FGR at different levels of circulating CORT (Sonnenberg et al. 2024), possibly through different sensitivity of feather follicles to CORT during moult (Lattin et al. 2012). Therefore, further research in how species differ in plasma CORT levels and CORT receptors during moult would be needed.

The lack of clear latitudinal signal in fCORT is in contrast with the observed pattern of lower circulating baseline breeding levels of CORT in tropical species (Hau et al. 2010, Kauzal et al. 2025; Table S5). But contrary to some tropical studies (Wada et al. 2006), we have observed a rise in circulating CORT outside the breeding season (but not necessarily moulting season) in our previous study on mostly the same Afrotropical species (Kauzal et al. 2025). This increase might be the reason why the difference in fCORT between tropical and temperate species in our dataset is smaller, thus more difficult to detect. In tropical species, fCORT could, on average, be biased towards higher values also by their more frequent moult breeding overlap (Moreno 2004). This overlap is probably rare, but when it occurs, it has been shown to lower the feather quality (Hemborg & Lundberg 1998, Echeverry-Galvis & Hau 2013) possibly due to high circulating CORT which is associated with breeding season in most species (Romero 2002). However, at least in the sampled Afrotropical species, the moult breeding overlap seems rather rare (Horák et al. 2022). Alternative explanation could be that CORT is simply downregulated to very similar levels in all species during moult, a period especially sensitive to elevated CORT (Romero 2002, Romero et al. 2005, Jenni-Eiermann et al. 2015). More research, especially sampling of circulating CORT during moult periods, is needed to disentangle these effects.

We did not find any support in our data for the association between fCORT and the migratory distance (Table S7). Even though migratory species often raise circulating CORT prior migration (Piersma et al. 2000, Eikenaar et al. 2014), most migratory passerines either strictly separate moult and migration in time, or suspend their moult for the migratory journey and resume it once they reach their target destination (Jenni & Winkler 2020a), effectively separating periods when high CORT is advantageous and when it is not. This might be especially important for migratory species in order to grow high quality feathers fast (De La Hera et al. 2009, 2012).

Among tropical birds, there was a tendency for species with higher elevational midpoints to show higher fCORT, similarly to a study conducted on lowland and highland populations of two temperate *Poecile* species (Sonnenberg et al. 2024). Authors of this study hypothesise that higher elevation populations are more time constrained by harsher climate and/or shorter moulting season (Kiat & Sapir 2021, Hernández=Téllez et al. 2024). We have found some support for a similar pattern across tropical species – fCORT being positively associated with species-specific elevational midpoint (though the credibility interval of this association also crossed zero: –0.01, 0.58; Table 3, Fig. 1). Lower temperatures of tropical mountains may select for higher CORT simply in order to compensate for potential energy losses at a time when plumage provides suboptimal thermoinsulation properties. High-elevation species may be therefore adapted to this increase in circulating CORT in order not to experience its detrimental effects on feather quality (Sonnenberg et al., 2024).

Another factor that can potentially influence the CORT content in feathers is their melanisation. Experimental work in pigeons suggests that darker parts of feathers (containing more eumelanin) bind CORT more strongly during growth, resulting in higher measured fCORT (Jenni-Eiermann et al. 2015). We did not measure melanin content in feathers (directly or via colour proxies) however, so we cannot test whether tropical and temperate species in our dataset differ systematically in tail-feather melanisation. Nevertheless, because whole plumage darkness covaries with humidity (more often higher in the tropics; Delhey et al. 2019), pigmentation differences could in principle add noise or even introduce a bias towards higher values in tropical species, when comparing fCORT across latitudes. While this mechanism alone is unlikely to be the only explanation for the lack of latitudinal contrast in our data, it remains unclear to what extent higher feather melanisation could obscure the lower corticosterone concentrations expected in tropical species, highlighting the need for future studies that integrate direct measures of feather pigmentation into comparative fCORT analyses.

To conclude, we were able to identify moderate within-species and high within-individual repeatability of fCORT and one potentially interesting trend regarding fCORT on a macroevolutionary scale – a positive association with altitude. Most notably, our data suggest that naturally occurring variation in CORT during moult (expressed as fCORT) is not the main driver of the previously reported differences in FGR and FBO between tropical and temperate zone passerines. The observed lack of clear latitudinal pattern in fCORT could suggest that seasonal downregulation of CORT during moult is indeed conserved adaptation in both temperate as well as in tropical species against detrimental effects of CORT on feather growth. We did, however, find a tentative support for higher fCORT of tropical spices breeding at higher elevations, consistent with increased overall energetical demands to moult in relatively harsher environment of tropical mountains. More generally, the high individual repeatability in our data indicate that fCORT is a robust “record” of circulating CORT during moult, and therefore can be informative in comparative or applied context like conservation biology. fCORT’s long term stability makes it especially useful to study non-invasively present and/or past stress individuals have experienced during moult, a critical period in life of most birds. To the best of our knowledge, this is a first comparative study on fCORT across diverse taxa and will hopefully spark future interest in this topic which will help clear some of the presented trends.

## Acknowledgements

We would like to thank to everyone who helped with the data collection in the field, namely Jaromír Čejka, Marek Brindzák, Kateřina Šimonová and Lukáš Bobek. Also, we are very grateful to the Chief of Bokwaongo, Chief of Bakingili, Francis Luma Ewome and the rest of Bokwaongo community for their help with the field work in Cameroon.

## Author contributions

OK and TA conceived and designed the study, all authors performed research, OK analysed the data, OK and TA drafted the manuscript with input from OT, and all authors contributed to revisions of the manuscript.

## Supplementary online material

### Supplementary Materials and methods

Detailed information about migration distance calculation, and information about the different methods employed to impute values of TEST below LOQ.

**Fig. S1**: Approximate locations of sampling localities (dotted white circles) with position of major town and cities (orange circles). Left: Czechia (temperate zone), right: Cameroon (tropical zone).

**Fig. S2:** Example tail of a typical passerine (Lesser Whitethroat, *Curruca curruca*) with individual rectrices numbered from inside out. Both second outermost tail feathers were collected for each individual bird if possible (“R5”, marked with arrows). For the analysis of fCORT the left feather was used in majority of cases (see main text).

**Fig. S3:** Relationship between fCORT_abs_ and species-specific body mass (left) and feather mass (right) showing the non-linear relationship on the original (absolute, not ln transformed) scale. The line on each plot represents the population-level relationship with a 95% credible interval and points show posterior median fitted values for each individual from the corresponding brms models.

**Table S1:** Number of fCORT samples below the limit of quantification (LOQ = 0.83 pg) out of the total number of samples for each group (BMC and sex). Percentages in parenthesis refer to the proportion of samples below LOQ for each group.

**Table S2:** Comparison of main models testing the association between fCORT and other variables using different methods to estimate values below LOQ (see electronic supplementary material, Materials and methods): i) imputing random values based on observed gaussian distribution, ii) assigning the value of LOQ/2 = 0.415 pg, and iii) using dataset without observations under LOQ (314 individuals of 86 species). Apart of phylogeny and run id, all models were controlled for species specific body mass, feather length and growth time, i.e. representing the differences in fCORT for same sized bird with same sized feather per one day.

All four models performed similarly. The most different model was the model using only values above the quantification limit with limited sample size. This is not surprising since the values below quantification limit were not randomly distributed but were rather more frequent in temperate species and especially so in temperate species moulting in tropics (Table S1).

**Table S3:** Summary of univariate models used to estimate the slope between fCORT, species specific body mass, feather mass and length. All models were controlled for phylogeny and in case of fCORT also for run id.

**Table S4:** Summary of predictor estimates of ASE models for fCORT using brms imputed values. Apart of phylogeny and run id, all models were controlled for species specific body mass, feather length and growth time, i.e. representing the differences in fCORT for same sized bird with same sized feather per one day.

**Table S5:** Summary of the main models testing the effect of breeding and moulting latitude (BMC) and sex on fCORT and their interaction (right model). Apart from phylogeny and run id, the models were controlled for species specific average body mass, feather length and growth time, i.e. representing the differences in fCORT for same sized bird with same sized feather per one day.

**Table S6:** Model comparison using the leave on out approach (*loo*) of different BRMS model testing the association of fCORT with BMC and sex. The “null” model did not contain any of these two. Apart from phylogeny and run id, the models were controlled for species specific average body mass, feather length and growth time.

**Table S7:** Summary of the model testing the effect of migration distance and sex on fCORT. Apart from phylogeny and run id, the model was controlled for species specific average body mass, feather length and growth time, i.e. representing the differences in fCORT for same sized bird with same sized feather per one day.

## Data availability

All data and scripts used to generate the results were uploaded to figshare and can be accessed here: https://doi.org/10.6084/m9.figshare.31017340.

