## Supplemental information for "Feather corticosterone is not associated with feather growth rate or quality across tropical and temperate passerines, but possibly linked to elevation"

Electronic Supplementary Material

**Journal:** Journal of Vertebrate Biology

**DOI:** xxx

### Supplementary Materials and methods

##### List of abbreviations

**ASE**: “adjusted single effect” model ­– model testing a single predictor but controlled for potentially confounding variables (see main text)

**BMC**: “breeding-moulting category”: variable combining breeding and moulting latitude of a bird species with three levels: i) temperate-temperate (reference level), ii) temperate-tropical and iii) tropical-tropical.

**FBO**: fault bar occurrence

**fCORT**: feather corticosterone

**fCORT_abs_**: absolute value of fCORT measured in a sample

**FGR**: feather growth rate

**LOQ**: limit of quantification

***pd***: probability of direction; measure of how much of the posterior mass was on one side of zero, which corresponds approximately to a two-sided frequentist *p-value* (*pd ≈ 1 – p value*; (Makowski *et al.* 2019)

##### Migration distance calculation

Migration distance was calculated as latitudinal arch difference (in 1000km) between the latitudinal centroid of breeding and wintering distribution of a species:

$$migration distance= \frac{\left( breeding latitude -wintering latitude \right)*Earth circumference}{360}*0.001$$

Breeding latitude and wintering latitude were in degrees (negative values for southern hemisphere); Earth circumference was set to 40 008 km (across poles). For breeding range centroid, we merged the “breeding” and “resident” range for a given species, for wintering range we merged the “wintering” and “resident” range for a given species from the original data (BirdLife International and Handbook of the Birds of the World 2018). All ranges were then restricted to the geographical limits of Europe (Western Palearctic sensu Shirihai & Svensson; 2018) and Africa, where all analysed species overwinter (Cepák *et al.* 2008). We made three exceptions to this rule:

- Data for Common Rosefinch (*Carpodacus erythrinus*) were not restricted, as unlike all the other analysed species, Common Rosefinch overwinters in Indian subcontinent (Lisovski *et al.* 2021).
- In order to distinguish the partially migratory European Stonechat (*Saxicola [torquatus] rubicola*) from the African Stonechat (*S. torquatus sensu stricto*; both treated as a single species *S. torquatus* in the distributional data), we restricted the breeding and wintering range of this (sub)species to the limits of Western Palearctic, which is a good enough approximation of its range (Shirihai & Svensson 2018).
- We used similar treatment for Eurasian reed-warbler (*Acrocephalus scirpaceus*) which is treated as a single species together with the sedentary African reed warbler (*Acrocephalus baeticatus*). Therefore, we restricted its breeding range to the limits of Western Palearctic and its wintering distribution by the limits of Afrotropical realm. This approach separates well the migratory and non-migratory subpopulations of these two species (Shirihai & Svensson 2018).

For all tropical breeding species, the migration distance was set to 0.

##### Values below limit of quantification (LOQ)

Due to the limited weight of the feather samples of small passerines (0.9–43.4 mg), 38 out of the 352 samples of fCORT measured were under quantification limit (LOQ = 0.83 pg in the final 60 μl extract (Adámková *et al.* 2019). Their distribution was non-random in the dataset and differed with respect to breeding and moulting latitude (see main text and Table S1). We employed several ways how to treat these values.

1) We used the *cens* function from the brms package (LOQ set to 0.83) to impute the missing data. This approach allows the Bayesian model to treat values below LOQ as left-censored data and estimate them as part of the posterior distribution accounting for their uncertainty (Bürkner 2017).
2) We estimated the mean and standard deviation of Gaussian distribution fitted to the ln-transformed data using *survreg* function from the package survival, which accounts for left censored data when estimating the distribution (Therneau 2025). We then generated random values from this distribution using *rnorm* function, excluding values above LOQ (set to ln (0.83)), and used these values in the models.
3) We replaced all values below LOQ with half the LOQ (0.415 pg), a commonly used approached for handling the left-censored data (Shoari & Dubé 2018).
4) We excluded all values below LOQ from the dataset and performed the models using only remaining observations reducing the dataset to 314 individuals of 86 species.

Despite using different imputation strategies for values below LOQ, all four main models performed similarly with comparable parameter estimates (c.f. Tables 1, 2 and 3 in main text and Table S2).

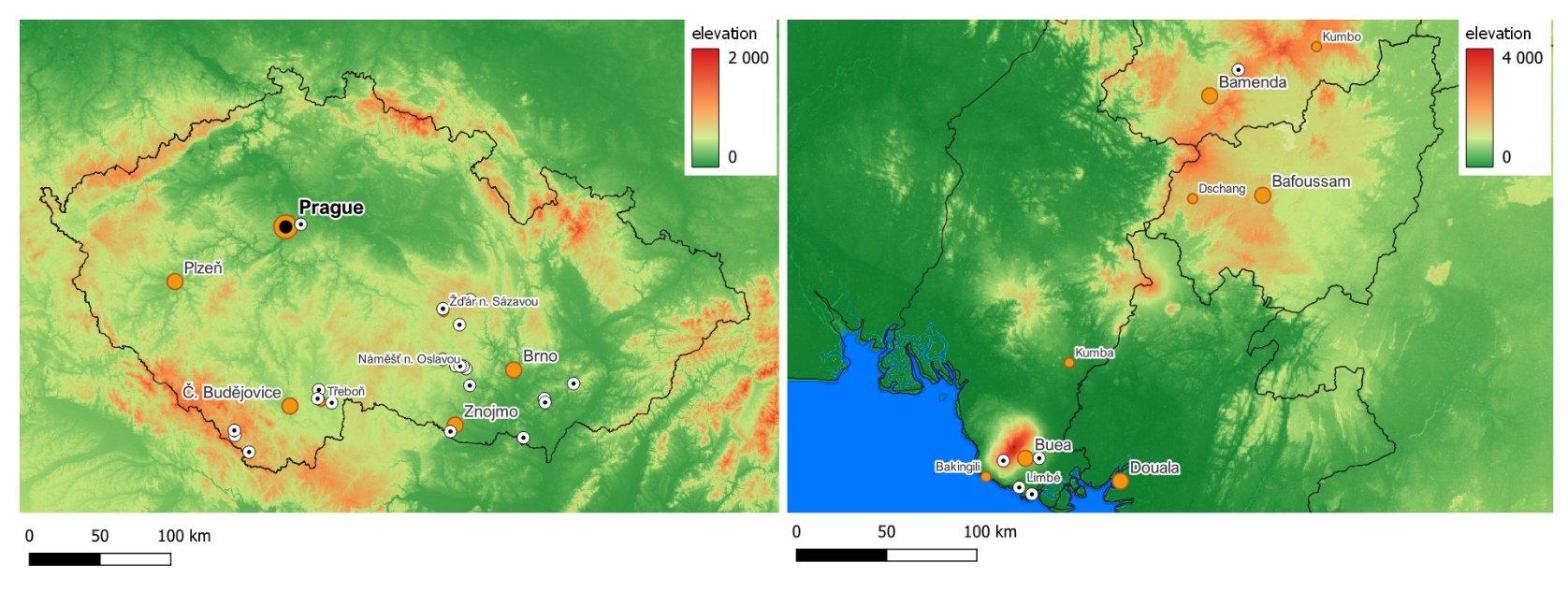

### Supplementary figures

**Figure S1:** Approximate locations of sampling localities (dotted white circles) with position of major town and cities (orange circles). Left: Czechia (temperate zone), right: Cameroon (tropical zone).

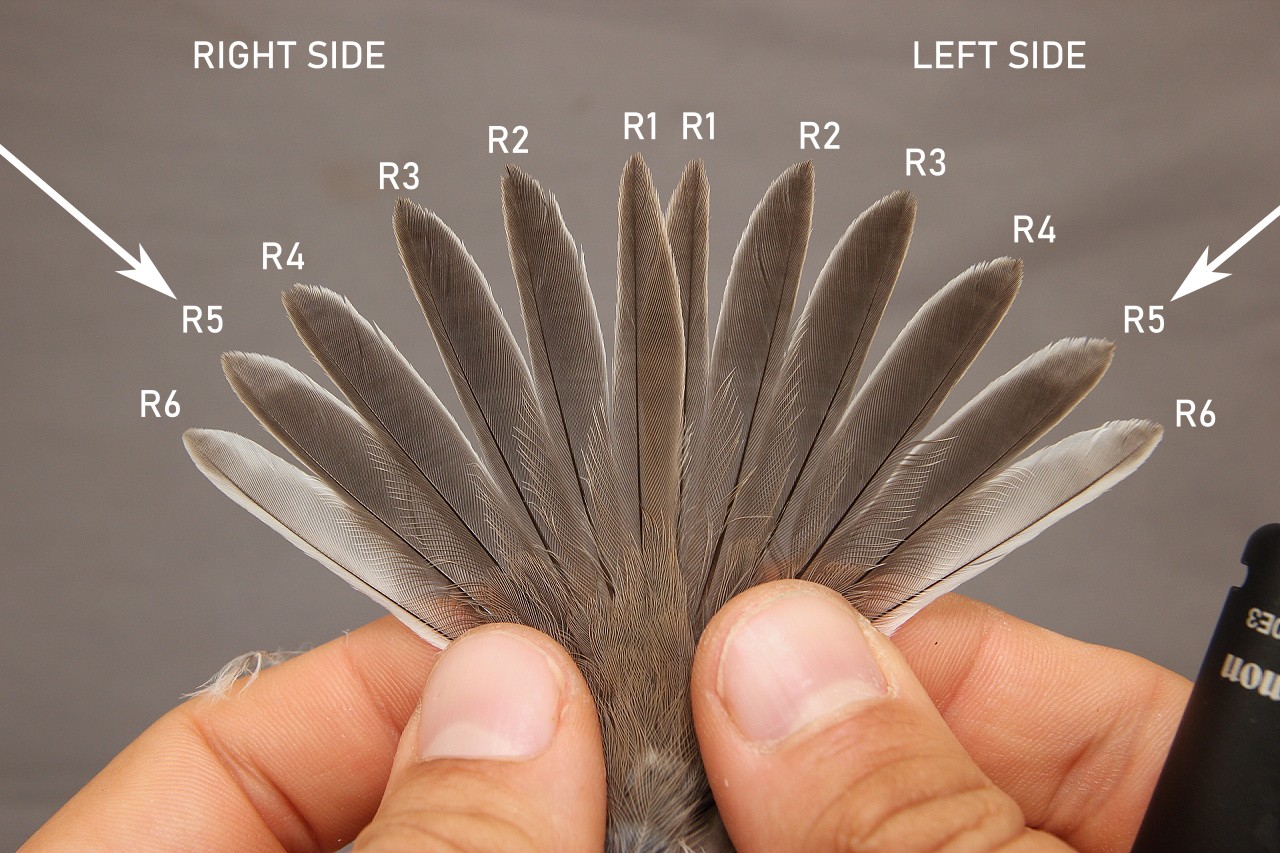

**Figure S2:** Example tail of a typical passerine (Lesser Whitethroat, *Curruca curruca*) with individual rectrices numbered from inside out. Both second outermost tail feathers were collected for each individual bird if possible (“R5”, marked with arrows). For the analysis of fCORT the left feather was used in majority of cases (see main text). *Photo: Ondřej Kauzál*.

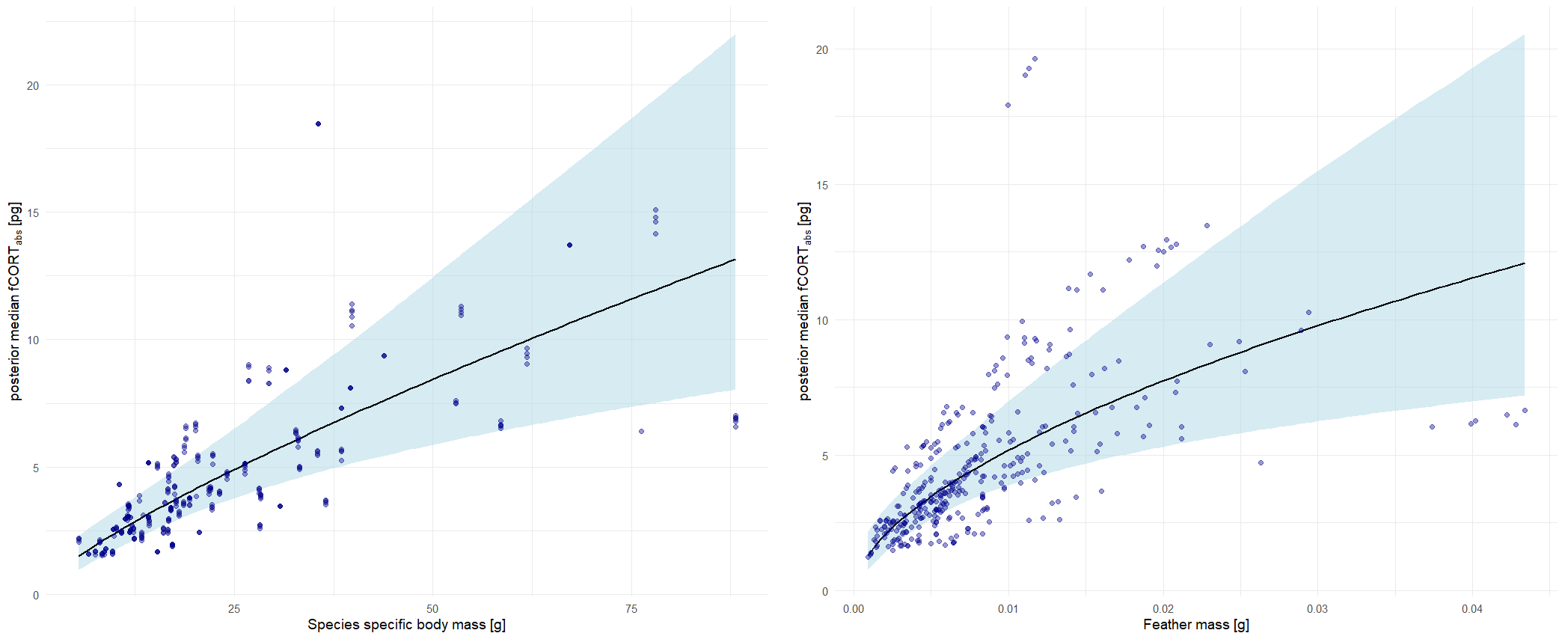

**Figure S3:** Relationship between fCORT_abs_ and species-specific body mass (left) and feather mass (right) showing the non-linear relationship on the original (absolute, not ln transformed) scale. The line on each plot represents the population-level relationship with a 95% credible interval and points show posterior median fitted values for each individual from the corresponding brms models. See also Table S3.

### Supplementary tables

**Table S1:** Number of fCORT samples below the limit of quantification (LOQ = 0.83 pg) out of the total number of samples for each group (BMC and sex). Percentages in parenthesis refer to the proportion of samples below LOQ for each group.

| **BMC** | **males** | **females** | *SUBTOTAL* |
| --- | --- | --- | --- |
| temperate-temperate | 10/78 | 6/49 | *16/127 (12.6 %)* |
| temperate-tropical | 7/37 | 4/15 | *11/46 (23.9 %)* |
| tropical-tropical | 6/90 | 5/89 | *11/179 (6.1 %)* |
| *SUBTOTAL* | *23/199 (11.6 %)* | *15/153 (9.8 %)* |  |

*Supplementary tables section continues on next page…*

**Table S2:** Comparison of main models testing the association between fCORT and other variables using different methods to estimate values below LOQ (see electronic supplementary material, Materials and methods): i) imputing random values based on observed gaussian distribution, ii) assigning the value of LOQ/2 = 0.415 pg, and iii) using dataset without observations under LOQ (*n* = 314 individuals of 86 species). For the FBO and FGR models where fCORT was a predictor, we substituted the values below LOQ with random values from estimated distribution in main text and here report only models where fCORT was imputed as LOQ/2 using dataset without observations under LOQ. Apart of phylogeny and run ID, all models were controlled for species specific body mass, feather length and growth time, i.e. representing the differences in fCORT for same sized bird with same sized feather per one day. Values with CrI95 not containing zero are highlighted in **bold** and regarded as significant support for an effect. Weakly supported trends in models (*pd* between 95% and 97.5%) are highlighted in *italic*.

All four models performed similarly. The most different model was the model using only values above the quantification limit with limited sample size. This is not surprising since the values below quantification limit were not randomly distributed but were rather more frequent in temperate species and especially so in temperate species moulting in tropics (Table S1).

a) Model testing the effect of BMC and sex on fCORT (for brms imputed values see Table S5):

| **predictor** | **random** | | | **LOQ/2 = 0.415 pg** | | | **only values > LOQ** (*n* = 314) | | |
| --- | --- | --- | --- | --- | --- | --- | --- | --- | --- |
|  | ***b*** | **LCrI** | **UCrI** | ***b*** | **LCrI** | **UCrI** | ***b*** | **LCrI** | **UCrI** |
| BMC: temperate–tropical | –0.33 | –0.89 | 0.22 | –0.38 | –0.94 | 0.17 | 0.08 | –0.41 | 0.57 |
| BMC: tropical–tropical | –0.17 | –0.56 | 0.22 | –0.15 | –0.55 | 0.26 | *–0.30* | *–0.64* | *0.04* |
| sex (female) | 0.16 | –0.07 | 0.39 | 0.13 | –0.11 | 0.37 | 0.12 | –0.06 | 0.31 |
| ln species specific body mass | **0.72** | **0.34** | **1.09** | **0.73** | **0.35** | **1.11** | **0.7** | **0.38** | **1.03** |
| ln feather length | –0.01 | –0.88 | 0.84 | –0.02 | –0.89 | 0.87 | 0.22 | –0.51 | 0.95 |
| ln growth time | 0.56 | –0.39 | 1.53 | 0.45 | –0.51 | 1.43 | 0.49 | –0.30 | 1.30 |
| Bayesian conditional R^2^ | 0.32 | 0.24 | 0.41 | 0.30 | 0.21 | 0.39 | 0.44 | 0.36 | 0.52 |
| Bayesian marginal R^2^ | 0.17 | 0.08 | 0.26 | 0.16 | 0.08 | 0.25 | 0.26 | 0.16 | 0.36 |

b) Model testing the effect of BMC and sex and its interaction on fCORT (for brms imputed values see Table S5):

| **predictor** | **random** | | | **LOQ/2 = 0.415 pg** | | | **only values > LOQ** (*n* = 314) | | |
| --- | --- | --- | --- | --- | --- | --- | --- | --- | --- |
|  | ***b*** | **LCrI** | **UCrI** | ***b*** | **LCrI** | **UCrI** | ***b*** | **LCrI** | **UCrI** |
| BMC: temperate–tropical | –0.18 | –0.81 | 0.44 | –0.21 | –0.84 | 0.42 | 0.30 | –0.25 | 0.85 |
| BMC: tropical–tropical | –0.04 | –0.49 | 0.41 | –0.01 | –0.47 | 0.45 | –0.15 | –0.53 | 0.24 |
| sex (female) | *0.37* | *–0.02* | *0.76* | *0.35* | *–0.05* | *0.75* | **0.37** | **0.06** | **0.68** |
| ln species specific body mass | **0.71** | **0.33** | **1.09** | **0.72** | **0.33** | **1.09** | **0.69** | **0.37** | **1.02** |
| ln feather length | 0.00 | –0.86 | 0.87 | 0.00 | –0.89 | 0.89 | 0.23 | –0.50 | 0.97 |
| ln growth time | 0.53 | –0.43 | 1.49 | 0.41 | –0.59 | 1.40 | 0.47 | –0.34 | 1.28 |
| sex (female) × temperate–tropical | –0.41 | –1.17 | 0.35 | –0.47 | –1.25 | 0.31 | *–0.59* | *–1.26* | *0.09* |
| sex (female) × tropical–tropical | –0.30 | –0.80 | 0.19 | –0.31 | –0.82 | 0.19 | *–0.34* | *–0.74* | *0.05* |
| Bayesian conditional R^2^ | 0.33 | 0.24 | 0.41 | 0.31 | 0.22 | 0.40 | 0.46 | 0.37 | 0.53 |
| Bayesian marginal R^2^ | 0.18 | 0.09 | 0.26 | 0.17 | 0.08 | 0.25 | 0.27 | 0.17 | 0.37 |

c) Model testing the effect of migration distance (controlled for breeding latitude) and sex on fCORT (for brms imputed values see Table S7):

| **predictor** | **random** | | | **LOQ/2 = 0.415 pg** | | | **only values > LOQ** (*n* = 314) | | |
| --- | --- | --- | --- | --- | --- | --- | --- | --- | --- |
|  | ***b*** | **LCrI** | **UCrI** | ***b*** | **LCrI** | **UCrI** | ***b*** | **LCrI** | **UCrI** |
| migration distance | –0.12 | –0.30 | 0.07 | –0.13 | –0.32 | 0.05 | 0.04 | –0.13 | 0.21 |
| sex (female) | 0.13 | –0.11 | 0.37 | 0.13 | –0.11 | 0.37 | 0.12 | –0.06 | 0.31 |
| breeding latitude (tropics) | –0.22 | –0.67 | 0.23 | –0.23 | –0.69 | 0.23 | –0.26 | –0.66 | 0.13 |
| ln species specific body mass | **0.68** | **0.30** | **1.05** | **0.69** | **0.31** | **1.07** | **0.71** | **0.39** | **1.05** |
| ln feather length | 0.03 | –0.83 | 0.89 | 0.01 | –0.87 | 0.88 | 0.23 | –0.52 | 0.97 |
| ln growth time | 0.34 | –0.63 | 1.32 | 0.44 | –0.56 | 1.43 | 0.49 | –0.32 | 1.29 |
| Bayesian conditional R^2^ | 0.29 | 0.20 | 0.38 | 0.30 | 0.21 | 0.39 | 0.44 | 0.36 | 0.52 |
| Bayesian marginal R^2^ | 0.15 | 0.07 | 0.24 | 0.16 | 0.08 | 0.24 | 0.26 | 0.16 | 0.36 |

d) Model testing the effect of the elevation and its deviation from mean and sex on fCORT (only tropical species, *n* = 172 ind. of 46 species; see main text Table 3 for brms imputed values):

| **predictor** | **random** | | | **LOQ/2 = 0.415 pg** | | | **only values > LOQ** (*n* = 168) | | |
| --- | --- | --- | --- | --- | --- | --- | --- | --- | --- |
|  | ***b*** | **LCrI** | **UCrI** | ***b*** | **LCrI** | **UCrI** | ***b*** | **LCrI** | **UCrI** |
| sex (female) | 0.07 | –0.20 | 0.34 | 0.05 | –0.24 | 0.32 | –0.01 | –0.23 | 0.22 |
| species elevation midpoint | *0.30* | *–0.01* | *0.60* | *0.28* | *–0.02* | *0.58* | **0.33** | **0.04** | **0.63** |
| diff. from species elevation midpoint | –0.15 | –0.50 | 0.21 | –0.15 | –0.50 | 0.22 | –0.20 | –0.53 | 0.12 |
| ln species specific body mass | **0.96** | **0.40** | **1.53** | **0.98** | **0.42** | **1.55** | **0.70** | **0.17** | **1.23** |
| ln feather length | –0.41 | –1.52 | 0.68 | –0.41 | –1.52 | 0.66 | 0.12 | –0.90 | 1.11 |
| ln growth time | 0.95 | –0.30 | 2.19 | 0.93 | –0.33 | 2.2 | 0.26 | –0.83 | 1.35 |
| Bayesian conditional R^2^ | 0.45 | 0.34 | 0.54 | 0.43 | 0.32 | 0.53 | 0.54 | 0.44 | 0.63 |
| Bayesian marginal R^2^ | 0.27 | 0.14 | 0.39 | 0.27 | 0.14 | 0.39 | 0.30 | 0.15 | 0.43 |

e) Model testing the effect of fCORT, BMC and sex on FGR; see main text Table 1 for values imputed from estimated distribution):

| **predictor** | **LOQ/2 = 0.415 pg** | | | **only values > LOQ** (*n* = 314) | | |
| --- | --- | --- | --- | --- | --- | --- |
|  | ***b*** | **LCrI** | **UCrI** | ***b*** | **LCrI** | **UCrI** |
| BMC: temperate–tropical | 0.14 | –0.10 | 0.38 | 0.13 | –0.13 | 0.38 |
| BMC: tropical–tropical | **–0.35** | **–0.51** | **–0.19** | **–0.36** | **–0.53** | **–0.19** |
| sex (female) | 0.00 | –0.06 | 0.06 | 0.00 | –0.06 | 0.07 |
| ln species specific fCORT_abs_ | 0.01 | –0.08 | 0.10 | 0.09 | –0.03 | 0.21 |
| fCORT_abs_ deviation from average | –0.02 | –0.05 | 0.01 | –0.02 | –0.06 | 0.01 |
| ln species specific body mass | **0.49** | **0.32** | **0.66** | **0.42** | **0.22** | **0.61** |
| Bayesian conditional R^2^ | 0.81 | 0.78 | 0.83 | 0.82 | 0.80 | 0.84 |
| Bayesian marginal R^2^ | 0.39 | 0.27 | 0.50 | 0.39 | 0.26 | 0.51 |

f) Model testing the effect of fCORT, BMC and sex on FBO; see main text Table 2 for values imputed from estimated distribution):

| **predictor** | **LOQ/2 = 0.415 pg** | | | **only values > LOQ** (*n* = 314) | | |
| --- | --- | --- | --- | --- | --- | --- |
|  | ***b*** | **LCrI** | **UCrI** | ***b*** | **LCrI** | **UCrI** |
| BMC: temperate–tropical | 0.46 | –1.83 | 2.55 | 1.16 | –1.29 | 3.51 |
| BMC: tropical–tropical | **2.62** | **1.39** | **4.07** | **2.84** | **1.44** | **4.60** |
| sex (female) | 0.11 | –0.59 | 0.81 | 0.16 | –0.59 | 0.92 |
| ln species specific fCORT_abs_ | –0.05 | –0.65 | 0.59 | –0.21 | –0.98 | 0.55 |
| fCORT_abs_ deviation from average | –0.17 | –0.57 | 0.23 | 0.00 | –0.44 | 0.43 |
| ln species specific body mass | 0.67 | –0.48 | 1.84 | 0.88 | –0.35 | 2.19 |
| ln feather length | –1.04 | –3.37 | 1.12 | –0.78 | –3.23 | 1.54 |
| ln growth time | 2.19 | –0.65 | 5.26 | 1.83 | –1.18 | 4.94 |
| Bayesian conditional R^2^ | 0.21 | 0.12 | 0.32 | 0.22 | 0.12 | 0.33 |
| Bayesian marginal R^2^ | 0.13 | 0.04 | 0.23 | 0.13 | 0.04 | 0.24 |

*Supplementary tables section continues on next page…*

**Table S3:** Summary of simple phylogenetic regression models used to estimate the slope between fCORT, species specific body mass, feather mass and length. All models were controlled for phylogeny and in case of fCORT also for run ID.

| **model** | ***b*** | **LCrI** | **UCrI** | **marginal *R^2^*** |
| --- | --- | --- | --- | --- |
| **ln fCORT_abs_ ⁓** |  |  |  |  |
| ln species specific body mass | 0.78 | 0.50 | 1.05 | 0.17 |
| ln feather mass | 0.58 | 0.34 | 0.81 | 0.13 |
| ln feather length | 0.94 | 0.31 | 1.55 | 0.06 |
| **ln feather mass ⁓** |  |  |  |  |
| ln species specific body mass | 0.99 | 0.85 | 1.12 | 0.73 |
| **ln feather length ⁓** |  |  |  |  |
| ln species specific body mass | 0.29 | 0.20 | 0.37 | 0.38 |

**Table S4:** Summary of predictor estimates of ASE models for fCORT using brms imputed values. Apart of phylogeny and run ID, all models were controlled for species specific body mass, feather length and growth time, i.e. representing the differences in fCORT for same sized bird with same sized feather per one day. Values with CrI95 not containing zero are highlighted in **bold** and regarded as significant support for an effect. Weakly supported trends in models (*pd* between 95% and 97.5%) are highlighted in *italic*.

| **predictor** | ***b*** | **LCrI** | **UCrI** | **marginal *R^2^*** |
| --- | --- | --- | --- | --- |
| breeding latitude (tropics) | –0.04 | –0.41 | 0.32 | 0.17 |
| moulting latitude (tropics) | –0.21 | –0.58 | 0.16 | 0.18 |
| BMC: temperate–tropical | –0.39 | –0.95 | 0.16 | 0.19 |
| BMC: tropical–tropical | –0.15 | –0.55 | 0.24 |  |
| sex (female) | 0.13 | –0.10 | 0.37 | 0.18 |
| FGR | –0.26 | –5.32 | 5.53 | 0.17 |
| FBO | –0.13 | –0.47 | 0.21 | 0.17 |
| temperate species only (171 individuals, 40 species): | | | | |
| *migration distance (1000km)* | *–0.20* | *–0.46* | *0.04* | 0.19 |
| tropical species only (174 individuals, 46 species): | | | | |
| **species elevation midpoint** | **0.32** | **0.04** | **0.60** | 0.28 |
| diff. from species elevation midpoint | –0.22 | –0.55 | 0.11 | 0.26 |

**Table S5:** Summary of the main models testing the effect of breeding and moulting latitude (BMC) and sex on fCORT and their interaction (right model). Apart from phylogeny and run ID, the models were controlled for species specific average body mass, feather length and growth time, i.e. representing the differences in fCORT for same sized bird with same sized feather per one day. Values with CrI95 not containing zero are highlighted in **bold** and regarded as significant support for an effect. Weakly supported trends in models (*pd* between 95% and 97.5%) are highlighted in *italic*.

| **predictor** | ***b*** | **LCrI** | **UCrI** | ***b*** | **LCrI** | **UCrI** |
| --- | --- | --- | --- | --- | --- | --- |
| BMC: temperate–tropical | –0.40 | –0.97 | 0.14 | –0.24 | –0.88 | 0.39 |
| BMC: tropical–tropical | –0.17 | –0.56 | 0.24 | –0.03 | –0.49 | 0.43 |
| sex (female) | 0.14 | –0.10 | 0.37 | *0.35* | *–0.04* | *0.74* |
| ln species specific body mass | **0.72** | **0.35** | **1.11** | **0.72** | **0.34** | **1.10** |
| ln feather length | –0.01 | –0.88 | 0.86 | 0.00 | –0.88 | 0.88 |
| ln growth time | 0.51 | –0.49 | 1.50 | 0.47 | –0.52 | 1.46 |
| sex (female) × temperate–tropical | – | – | – | –0.45 | –1.24 | 0.33 |
| sex (female) × tropical–tropical | – | – | – | –0.31 | –0.81 | 0.19 |
| effect of phylogeny | 0.19 | 0.00 | 0.44 | 0.18 | 0.00 | 0.44 |
| Bayesian conditional R^2^ | 0.36 | 0.27 | 0.45 | 0.37 | 0.28 | 0.46 |
| Bayesian marginal R^2^ | 0.20 | 0.10 | 0.30 | 0.21 | 0.11 | 0.30 |

**Table S6:** Model comparison using the leave on out approach (*loo*) of different BRMS model testing the association of fCORT with BMC and sex. The “null” model did not contain any of these two. Apart from phylogeny and run ID, the models were controlled for species specific average body mass, feather length and growth time. For detailed summary of the first two models see Table S5.

| **predictor** | ***Δelpdloo*** | ***s.d.* of the difference** |
| --- | --- | --- |
| BMC × sex | –2.3 | 2.6 |
| BMC + sex | –0.9 | 1.9 |
| BMC | –0.6 | 1.3 |
| sex | –0.3 | 1.2 |
| “NULL” | 0 | 0 |

**Table S7:** Summary of the model testing the effect of migration distance and sex on fCORT on a subset of temperate species (171 individuals, 40 species). Apart from phylogeny and run ID the model was controlled for species specific average body mass, feather length and growth time, i.e. representing the differences in fCORT for same sized bird with same sized feather per one day. Values with CrI95 not containing zero are highlighted in **bold** and regarded as significant support for an effect. Weakly supported trends in models (*pd* between 95% and 97.5%) are highlighted in *italic*.

| **predictor** | ***b*** | **LCrI** | **UCrI** |
| --- | --- | --- | --- |
| migration distance | –0.19 | –0.45 | 0.05 |
| sex (female) | 0.20 | –0.21 | 0.62 |
| *ln species specific body mass* | *0.62* | *–0.01* | *1.23* |
| ln feather length | 0.07 | –1.78 | 1.97 |
| ln growth time | –0.28 | –1.94 | 1.38 |
| effect of phylogeny | 0.24 | 0.00 | 0.55 |
| Bayesian conditional R^2^ | 0.34 | 0.20 | 0.45 |
| Bayesian marginal R^2^ | 0.20 | 0.07 | 0.34 |
